# Bone marrow lymphocyte dynamics during chemotherapy in pediatric acute myeloid leukemia

**DOI:** 10.1101/2025.05.04.652110

**Authors:** Joost B. Koedijk, Farnaz Barneh, Joyce E. Meesters-Ensing, Marc van Tuil, Edwin Sonneveld, Sander Lambo, Alicia Perzolli, Elizabeth K. Schweighart, Mauricio N. Ferrao Blanco, Merel van der Meulen, Anna Deli, Elize Haasjes, Kristina Bang Christensen, Hester A. de Groot-Kruseman, Soheil Meshinchi, Henrik Hasle, Mirjam E. Belderbos, Maaike Luesink, Bianca F. Goemans, Stefan Nierkens, Jayne Hehir-Kwa, C. Michel Zwaan, Olaf Heidenreich

## Abstract

A better understanding of lymphocyte dynamics during current treatment regimens in pediatric AML is urgently needed to understand whether the application of bispecific T-cell-engagers (BiTEs) during periods of low tumor burden could be a viable treatment strategy. In this study, we found that induction 1, comprising mitoxantrone-etoposide-cytarabine in nearly all patients (as part of the NOPHO-DBH AML-2012 protocol), led to preserved or increased relative lymphocyte abundances alongside marked blast reduction in most cases. This was accompanied by a shift towards higher T-cell fractions, potentially creating a favorable window for BiTE therapy. The absence of a correlation between blast reduction and lymphocyte changes suggests that chemotherapy exerts differential effects on the lymphocyte compartment. Despite the heterogeneity of agents used in induction 2, more than half of patients showed a decline in lymphocyte levels. Nonetheless, the increase in T- and B-cells observed in most patients from the NOPHO-AML 2004 cohort after induction 2 suggests that lymphocyte recovery at this treatment stage is not uniformly impaired. Our transcriptomic and ex vivo functional data align with preclinical findings in adult AML and provide a basis for further investigations in in vivo models and early clinical trials. Such efforts should prioritize novel BiTE constructs targeting multiple tumor-associated (e.g., NCT05673057) or tumor-specific antigens.

---

T-cell-directed immunotherapy, which aims to boost or induce T-cell-mediated anti-tumor immunity, has shown remarkable success in various cancers, including B-cell precursor acute lymphoblastic leukemia (BCP-ALL), making it a compelling avenue for investigation in acute myeloid leukemia (AML)^1, 2^. Bispecific T-cell-engagers (BiTEs) are a promising form of T-cell-directed immunotherapy that redirect CD3^+^ T-cells to tumor cells, thereby inducing T-cell activation and subsequent tumor cell lysis^3^. However, BiTEs, mainly targeting CD33 or CD123, have shown limited efficacy and/or high toxicity in relapsed/refractory AML^4-7^. A proposed strategy to enhance BiTE therapy in AML is their administration during periods of minimal residual disease, e.g., between chemotherapy courses, as demonstrated in BCP-ALL^2, 8-10^. Chemotherapy may, however, significantly alter the immune landscape^11^: anthracyclines, for example, can promote anti-tumor immunity via immunogenic cell death^12^, but chemotherapy may also deplete lymphocytes and induce T-cell dysfunction^13, 14^. Since pre-treatment T-cell infiltration and dysfunction in the tumor microenvironment are key predictors of BiTE efficacy^15-18^, understanding how chemotherapy alters the immune landscape in the leukemic bone marrow (BM) is crucial for assessing the potential of BiTEs in between chemotherapy courses in AML. Given differences in disease biology, immune system maturity, and treatment regimens between pediatric and adult AML^19, 20^, pediatric-specific studies are necessary. Here, we examined the impact of chemotherapy-based regimens on the BM lymphocyte compartment in newly diagnosed pediatric AML (pAML).

We first characterized the treatment-naïve pAML BM lymphocyte compartment using diagnostic bulk RNA-sequencing (RNA-seq) data (**Fig.1A**). To reliably infer the lymphocyte composition from bulk RNA-seq data, we acquired a publicly-available single cell (sc) RNA-seq dataset^21^ to generate a healthy BM cell type signature matrix for use with CIBERSORTx^22^. To validate its performance, we retrieved BM scRNA-seq data from 27 pAML cases at diagnosis, remission, and/or relapse^23^, and generated pseudo-bulk profiles (n=62). Applying CIBERSORTx with the healthy BM reference to these pseudo-bulk profiles and comparing the deconvoluted estimates with the original scRNA-seq annotations (**Fig.S1A**), we observed strong correlations for T-, B- and NK-cells (T-cells: *r*=0.72, P<0.001; B-cells: *r*=0.87, P<0.001; NK-cells: *r=*0.68, P<0.001; **Fig.1B**). Similarly, CD4^+^ naïve, CD8^+^ effector, and CD8^+^ memory T-cells showed good concordance, while CD4^+^ memory and CD8^+^ naïve T-cells did not (**Fig.S1B**), supporting the method’s accuracy for most but not all lymphocyte subsets. Applying this approach to our primary study cohort (51 newly diagnosed pAML cases and seven age-matched controls; **Fig.1A**; **Table S1**), we found significantly lower fractions of T- and B-cells in the pAML BM compared to controls (P<0.001 and P=0.012, respectively; **Fig.1C**). Specifically, CD4^+^ naïve, CD8^+^ effector, and CD8^+^ memory T-cells were all less abundant (**Fig.S1C**). NK-cell fractions did not differ (**Fig.1C**). These findings, as anticipated, indicate a diminished lymphocyte compartment in the BM in newly diagnosed pAML.

**Figure 1.**
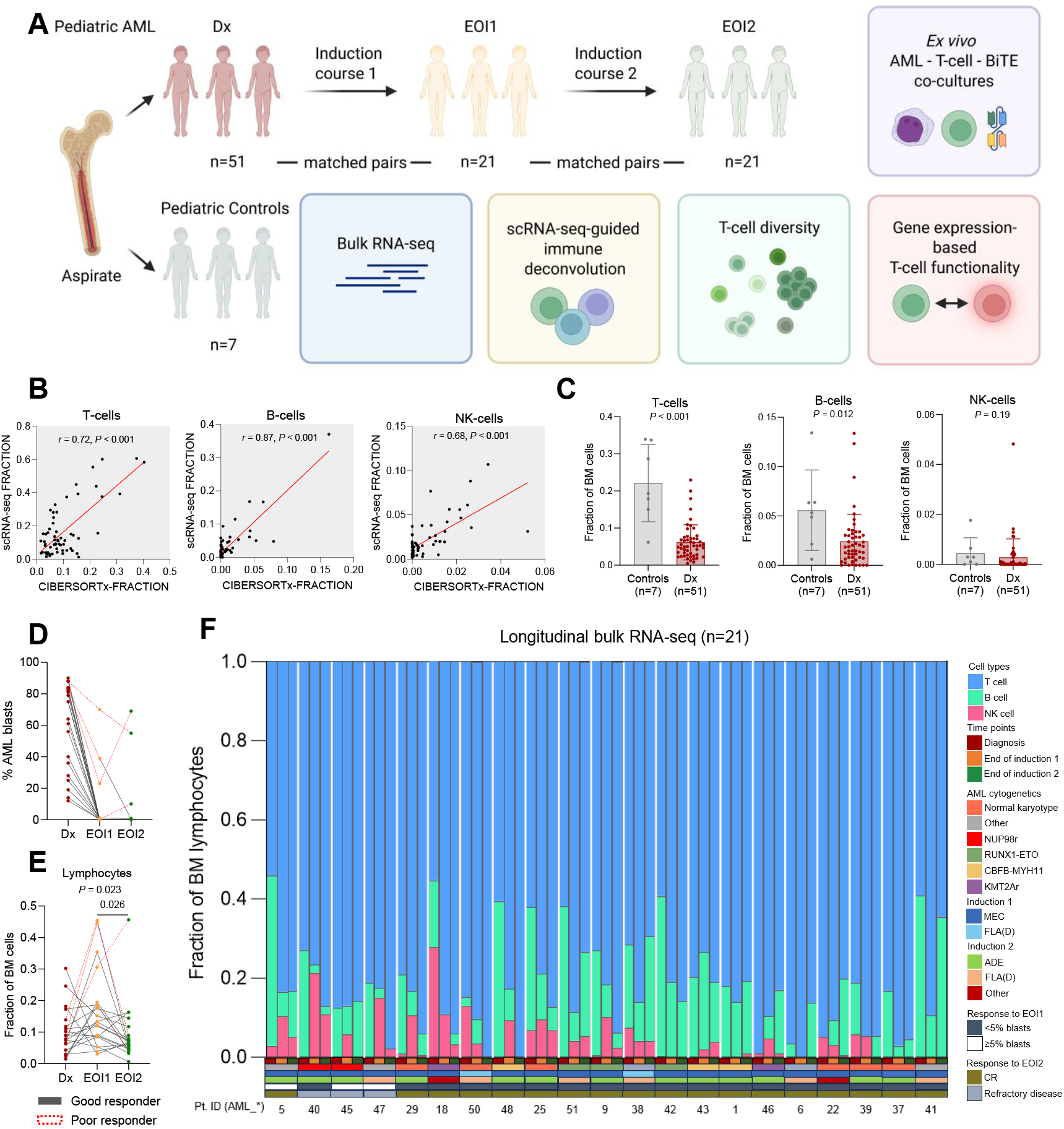
Study design and bone marrow lymphocyte dynamics in the primary study cohort. (A) Schematic overview of the study population and employed methods. Dx: diagnosis; EOI1: end of induction 1; EOI2: end of induction 2; bulk RNA-seq: bulk RNA-sequencing; scRNA-seq: single-cell RNA-sequencing; BiTE: bispecific T-cell engager. (B) Correlation plot of the positive correlation between the scRNA-seq- and CIBERSORTx-based fractions (relative abundances of cell populations out of all BM cells) of T-, B-, and natural killer (NK)-cells, calculated using Pearson correlation. (C) Comparison of the CIBERSORTx-based fractions of T-, B-, and NK-cells in the bone marrow (BM) of newly diagnosed pediatric AML patients at diagnosis versus non-leukemic controls (Mann-Whitney test). (D) Changes in AML blasts (measured by flow cytometry) over the course of treatment. (E) Comparison of CIBERSORTx-based lymphocyte abundance, out of BM cells, throughout induction therapy (Friedman test followed by Dunn’s multiple comparison test; the top value corresponds to the Friedman test, and the lower values indicate Dunn’s test results). (F) Overview of the CIBERSORTx-based fractions of T-, B-, and NK-cells out of BM lymphocytes for the 21 patients with longitudinal bulk RNA-seq data per treatment time point. The cytogenetic alterations in the ‘Other’ cytogenetic category are described for each patient in **Table S1**. MEC: mitoxantrone, etoposide, cytarabine; ADE: cytarabine, daunorubicin, etoposide; FLA(D): fludarabine, cytarabine with or without daunorubicin. The specific chemotherapeutic treatments used in the ‘Other’ category are provided in **Table S1**. Refractory disease was defined as the presence of ≥5% leukemic cells in the BM after the second induction course. CR: complete remission.

To investigate BM lymphocyte dynamics during chemotherapy, we performed bulk RNA-seq on 42 BM samples from 21 pAML cases, collected at end of induction 1 (EOI1) and EOI2 (similar time intervals, P=0.62, **Fig. S1D**). All patients were treated according to the NOPHO-DBH AML-2012 protocol (**Fig.1A,F**; **Table S1**). During induction 1, 19/21 patients received mitoxantrone, etoposide, and cytarabine (MEC). At EOI1, eighteen patients had good responses (<5% blasts by flow cytometry), while three (AML5, AML45, AML47) were poor responders (**Fig.1D,F**). Among good responders, lymphocyte fractions increased (>125% of baseline) in ten patients, remained stable (75-125%) in four, and decreased (<75%) in four (**Fig.1E-F**). Although an increase in lymphocyte fraction was expected due to the substantial blast clearance (median 62.5% to 0.1%), lymphocyte changes did not correlate with blast reduction (*r*=-0.32, P=0.20; n=18; **Fig.S2A**), suggesting differential effects of MEC on the BM lymphocyte compartment. No specific cytogenetic alterations were associated with a particular direction of lymphocyte change, which was expected due to the relatively small number of cases. Notably, all three poor responders showed marked lymphocyte increases at EOI1 (median 392%, range: 327-492%), despite high residual AML burden (median 39%, range: 23-70%; **Fig.1E**), suggesting that significant lymphocyte infiltration and/or expansion can occur even in the context of persistent leukemic infiltration. Lymphocyte subset analysis revealed that T-cells predominated at diagnosis (mean 75±21%) and further increased by EOI1 (mean 86±6.5%, P=0.016; **Fig.1F, Fig.S2B**). Within the T-cell compartment, CD4^+^ naïve T cells represented the most abundant subset at diagnosis (mean 57±26%), followed by CD8^+^ memory (29±14%) and CD8^+^ effector T-cells (5.8±6.9%, **Fig.S2C-D**). CD4^+^ naïve and CD8^+^ memory T-cell proportions remained largely stable following induction 1 (63±11%, P=0.68 and 21±10%, P=0.30, respectively), whereas CD8^+^ effector T-cells increased (11±5.2%, P=0.004; **Fig.S2C-D**). B-cell fractions decreased from 21±14% at diagnosis to 7.5±6.1% at EOI1 (P=0.006), while NK-cell proportions varied without consistent directional change (3.4±6.6% vs. 6.1±5.6%; P=0.37; **Fig.S2B**). Altogether, following induction 1, most patients showed increased or stable lymphocyte proportions alongside marked blast reduction. This was accompanied by a shift in the lymphoid compartment towards a higher T-cell fraction – in particular CD8^+^ effector T-cells - whereas B-cell proportions declined.

During induction 2, chemotherapy regimens were more heterogeneous: thirteen patients received ADE (cytarabine, daunorubicin, and etoposide), five FLA(D) (fludarabine and cytarabine ± daunorubicin), and three other regimens (**Fig.1F**; **Table S1**). Seventeen patients maintained remission, whereas one (AML40) showed disease progression (from 0.3 to 10%; **Fig.1D,F**). Of the three initial poor responders, AML5 achieved remission, whereas AML45 and AML47 had persistent disease (>5%; **Fig.1D,F**). Despite regimen variability, lymphocyte fractions declined significantly at EOI2 (P=0.026; **Fig.1E**). No differences between ADE and FLA(D)-treated patients were observed, though small group sizes precluded statistical testing (**Fig. S2E**). The abundance of T-cells out of total lymphocytes remained stable in most cases, while B-cell fractions frequently increased (eleven >125%, four 75-125%, six <75%) and NK-cell levels declined in nearly two-thirds of patients (13/21, P=0.09; **Fig.S2B**). Taken together, despite diverse treatment regimens, more than half of patients experienced a decline in total lymphocyte levels following induction 2, contrasting with the earlier induction phase.

To assess whether induction therapy was associated with changes in T-cell diversity, we profiled the T-cell receptor (TCR) repertoire using MiXCR^24^ (successful in 20/21 cases; **Fig.2A**). Shannon diversity indices increased from diagnosis to EOI1 and EOI2 (P=0.008 and P=0.08, respectively), but remained within a relatively narrow range throughout induction therapy in most patients (EOI1: 75-125% in 15/20 patients, >125% in 4, <75% in 1; EOI2: 75-125% in 17, >125% in 2, <75% in 1), indicating only modest changes in overall TCR diversity (**Fig. 2B**). Identical CDR3 β-chain sequences were detected at multiple timepoints in 8/20 cases, representing a median of 1.9% of the repertoire (range 0.3–5.1%; **Fig.2C**). These data suggest that chemotherapy is associated with a diverse and largely distinct post-treatment T-cell repertoire. Whether this includes tumor-reactive clones remains unclear. Future studies should investigate the tumor-specificity of T-cells persisting or emerging during therapy, as these may enhance BiTE efficacy^25^. In addition, the modest sensitivity of bulk RNA-seq-based TCR repertoire profiling requires validation using dedicated TCR-sequencing approaches.

**Figure 2.**
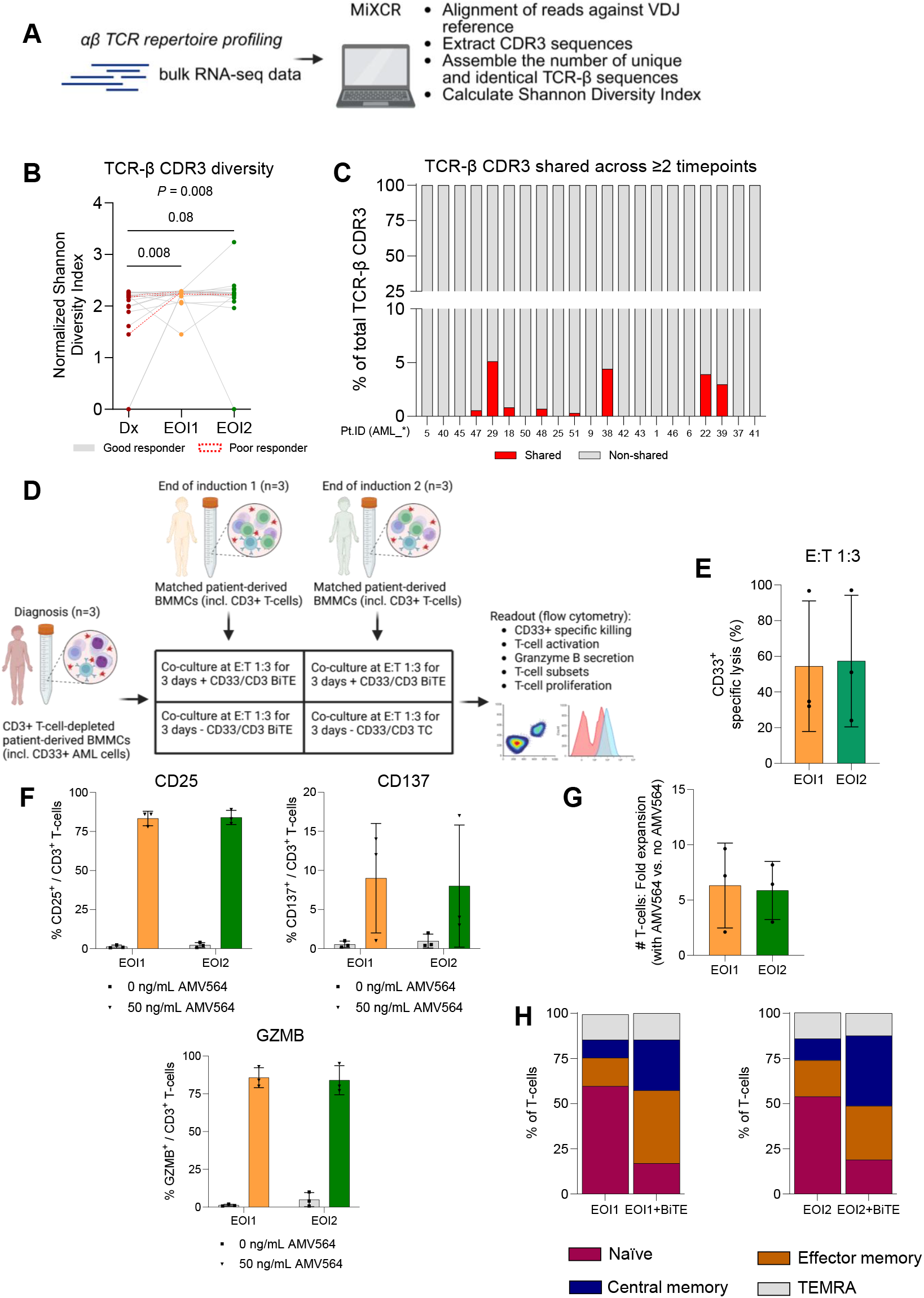
T-cell receptor repertoire profiling using bulk RNA-sequencing and *ex vivo* assays. (A) Schematic overview of the approach for estimating the αβ T-cell diversity from bulk RNA-sequencing using the MiXCR method. TCR: T-cell receptor; bulk RNA-seq: bulk RNA-sequencing; CDR3: complementary determining region 3. (B) Comparison of the normalized Shannon Diversity Index at diagnosis (Dx), end of induction 1 (EOI1), and end of induction 2 (EOI2; Friedman test followed by Dunn’s multiple comparison test; the top value corresponds to the Friedman test, and the lower values indicate Dunn’s test results). (C) Overview of the fraction of TCR-β CDR3 sequences that are shared between ≥2 timepoints for each patient. (D) Schematic overview of the experimental setup of the *ex vivo* assays. BMMCs: BM mononuclear cells; E:T: effector-to-target; BiTE: bispecific T-cell-engager. (E) CD33^+^ specific killing at EOI1 and EOI2 (normalized to CD33^+^ cell numbers in the absence of AMV564 (CD33/CD3-BiTE)). Dots indicate biological replicates (n=3 patients). (F) CD25-/CD137-/Granzyme B (GZMB)-positive CD3^+^ T-cells relative to all CD3^+^ T-cells at EOI1 and EOI2 after co-culture in the presence or absence of AMV564. Squares/triangles indicate biological replicates (n=3 patients). (G) T-cell numbers in the presence or absence of AMV564, depicted as fold expansion of the one (presence of AMV564) versus the other (absence of AMV564) condition, at EOI1 and EOI2. (H) Fraction of naive (CD45RA^+^CCR7^+^), effector memory (CD45RA^-^CCR7^-^), central memory (CD45RA^-^CCR7^+^), and terminally differentiated (TEMRA: CD45RA^+^CCR7^-^) T-cells relative to all CD3^+^ T-cells at EOI1 and EOI2 in the absence or presence of the BiTE AMV564.

Given its relevance for responses to T-cell-directed immunotherapies, we next assessed T-cell functionality^15, 16^. To this end, we applied established gene signature scores for T-cell cytolytic activity^26^, exhaustion^27^, and senescence^28^ to our bulk RNA-seq dataset, normalized for T-cell abundance. Cytolytic scores increased significantly following induction 1 (P=0.001) and remained elevated at EOI2 (P=0.002), whereas senescence and exhaustion scores varied considerably between patients without consistent directional change (**Fig.S2F**). To further evaluate the functionality of T-cells in between chemotherapy courses in pAML, we next investigated the ability of a CD33/CD3-BiTE (AMV564) to induce AML cell lysis via autologous T-cells derived from EOI1 or EOI2 BM mononuclear cells (BMMCs). Co-culturing EOI1/EOI2 BMMCs with CD3^+^ T-cell-depleted diagnostic BMMCs containing CD33^+^ AML cells (effector-to-target ratio 1:3) for three days in the presence or absence of AMV564 showed robust CD33^+^ cell lysis (mean specific lysis: 54±37% at EOI1, 57±37% at EOI2; **Fig.2D-E**). AMV564-induced cytotoxicity was accompanied by robust T-cell activation, evidenced by the upregulation of the T-cell activation markers CD25 and CD137, granzyme B expression, and T-cell proliferation (**Fig.2F-G**). Subset analysis revealed that BiTE therapy led to a phenotypic shift of naive to effector memory and central memory T-cells (**Fig.2H**). These data suggest that cytolytic potential increases, and that BiTE treatment is capable of activating autologous T-cells *ex vivo* and inducing lysis of primary CD33^+^ cells, at timepoints between chemotherapy cycles. While these results indicate functional T-cell potential at EOI1 and EOI2, further studies are required to determine consistency, T-cell function relative to healthy donor T-cells, and *in vivo* relevance.

Finally, we assessed BM lymphocyte dynamics in patients treated on other pAML protocols. Using data from the COG AAML1031 protocol (BM scRNA-seq data from seven treatment-naïve patients with paired diagnosis-EOI1 samples^23^; **Table S2**) and NOPHO-AML 2004 protocol (immunohistochemistry data for thirteen patients with diagnosis-EOI1-EOI2 BM samples^29^; **Table S3**), we observed both similarities and discrepancies in lymphocyte dynamics compared to the primary study cohort. Consistent with our previous findings, six out of seven COG patients (cytarabine, daunorubicin, etoposide, bortezomib/sorafenib) showed increased lymphocyte levels at EOI1 (P=0.031; **Fig.S3A-C**). Furthermore, the proportion of T-cells out of lymphocytes increased, while B-cells declined, and NK-cell changes were variable (**Fig. S3B**). Conversely, NOPHO-AML 2004 patients (course 1: cytarabine, idarubicin, etoposide, 6-thioguanine; course 2: cytarabine, mitoxantrone) exhibited profound T-cell heterogeneity and a reduction in B-cells at EOI1 (P=0.003; **Fig.S3E-F;** considering T- and B-cells in aggregate was not feasible because of the single-stain IHC). By EOI2, B-cell proportions recovered (P=0.007; **Fig.S3F**) and nine out of thirteen patients showed increased (>125%) T-cell levels, which was notable given the shorter EOI1–EOI2 interval compared to the NOPHO-DBH AML-2012 protocol (P<0.001; **Fig.S3G**). These findings suggest common trends in BM lymphocyte dynamics but also highlight protocol-specific variations, possibly linked to the use of specific chemotherapeutic agents. Given the limited number of patients in these external pAML cohorts, confirmation in larger cohorts is warranted.

A better understanding of lymphocyte dynamics during current treatment regimens in pAML is urgently needed to understand whether the application of BiTEs during periods of low tumor burden could be a viable treatment strategy. In this study, we found that induction 1, comprising MEC in nearly all patients, led to preserved or increased relative lymphocyte abundances alongside marked blast reduction in most cases. This was accompanied by a shift towards higher T-cell fractions, potentially creating a favorable window for BiTE therapy^15, 18^. The absence of a correlation between blast reduction and lymphocyte changes suggests that chemotherapy exerts differential effects on the lymphocyte compartment. Further studies are needed to clarify the mechanisms underlying divergent lymphocyte recovery, which may support the adaptation of treatment regimens to optimize conditions for immunotherapeutic interventions. Despite the heterogeneity of agents used in induction 2, more than half of patients showed a decline in lymphocyte levels. Nonetheless, the increase in T- and B-cells observed in most patients from the NOPHO-AML 2004 cohort after induction 2 suggests that lymphocyte recovery at this treatment stage is not uniformly impaired. Our transcriptomic and *ex vivo* functional data align with preclinical findings in adult AML^30^ and provide a basis for further investigations in *in vivo* models and early clinical trials. Such efforts should prioritize novel BiTE constructs targeting multiple tumor-associated (e.g., NCT05673057) or tumor-specific antigens.

## Author contributions

Conceptualization: JBK, SN, CMZ, OH. Methodology: JBK, FB, JEM-E, MT, ES, SL, AP, EKS, MNFB, BG, SN, CMZ, OH. Data acquisition: JBK, FB, JEM-E, MT, ES, SL, AP, EKS, EH, KBC, HAGK, SM, HH, MEB, BG, ML, JH-K, CMZ. Data analysis and interpretation: JBK, FB, JEM-E, SL, AP, EKS, MNFB, MM, AD. Writing – Original Draft: JBK. Writing – Reviewing & Editing: all authors. Supervision: SN, CMZ, OH.

## Acknowledgements

We would like to thank all patients and/or their families for their generous consent for the research use of these samples; the staff of the University Medical Center Utrecht Tissue Facility (Domenico Castigliego, Petra van der Weide, Petra van der Kraak, Erica Siera, Karina Timmer, Sven van Kempen); the MIMIC AML study team (Monique Schippers, Marijn Scheijde-Vermeulen, Gertjan Kaspers, Katja Heitink-Pollé, Arjenne Kors, Marleen Winnubst-Wassens, Dominique Mangert, Corinne Gerhardt, Margot Geerdink, Yvonne Ruchti, Lettine van den Brink, Ria Koolma, Nienke Adriaens, Danny Baars); the MIMIC Brain study team (Jasper van der Lugt, Friso Calkoen, Raoull Hoogendijk); the Princess Máxima Center biobank and diagnostic lab staff (Jantien Woudstra, Marion Persoon, Patty Arikan-Kok, Anja de Jong, Lisa Jacobi, Rubina Moeniralam, Arie Maat); Jeroen van Velzen (flow cytometry facility); the members of the Heidenreich group and Rico Hagelaar (Van Boxtel group). We apologize to authors whose papers we could not cite due to space limitations. We used ChatGPT (OpenAI) for assistance with language editing, focusing on grammar, phrasing, and consistency, under the authors’ supervision.

## Supplementary Figure Legends

**Supplementary Figure 1.**
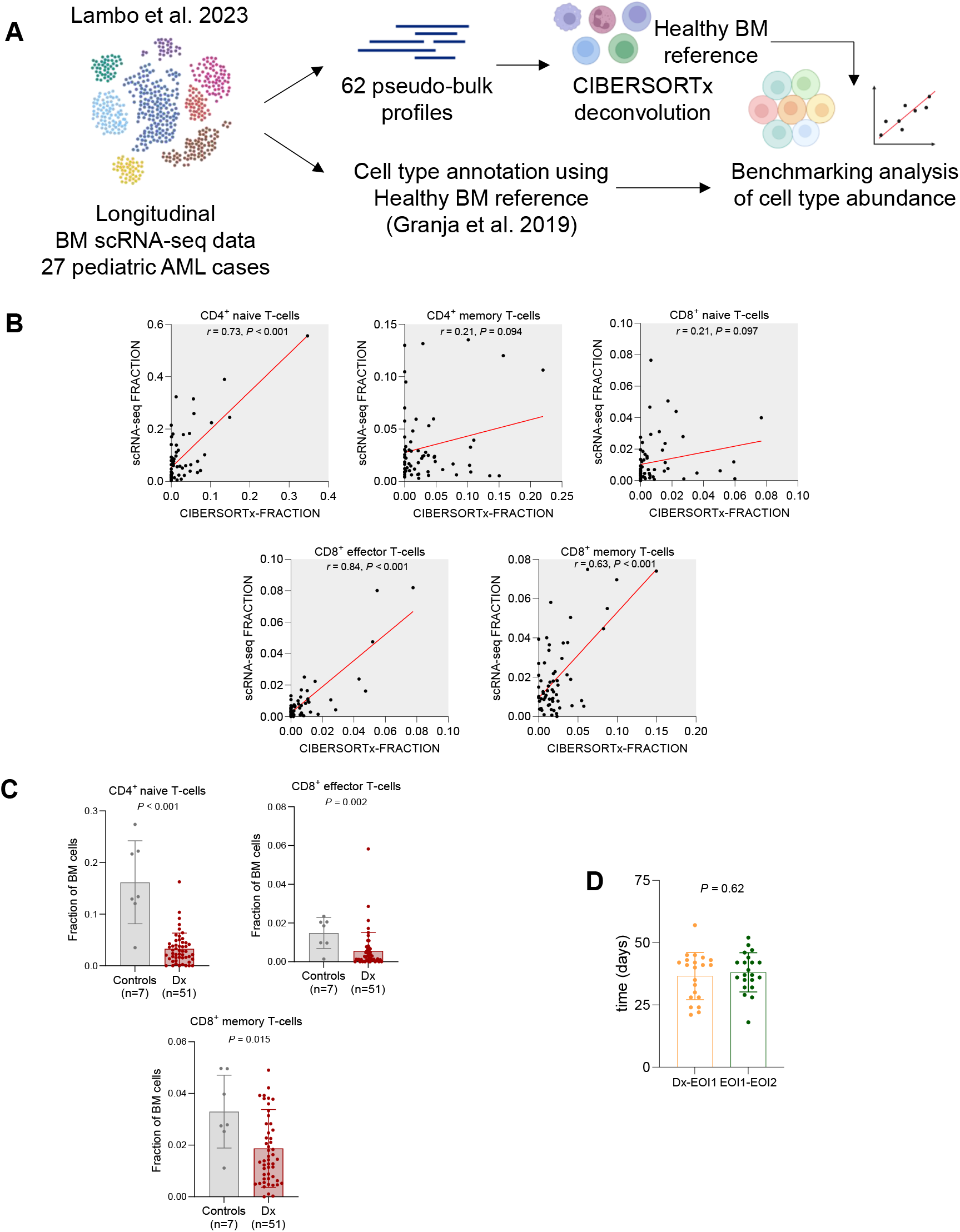
Benchmarking of single-cell RNA-sequencing-based immune deconvolution and its application in pediatric AML patients and non-leukemic controls. (A) Schematic overview of the benchmarking of the single-cell RNA-sequencing (scRNA-seq)-based immune deconvolution approach. BM: bone marrow. (B) Correlation plots showing the relationship between the scRNA-seq- and CIBERSORTx-based fractions (relative abundances) of T-cell subsets, calculated using Pearson correlation. (C) Comparison of the CIBERSORTx-based fraction (relative abundance) of validated T-cell subsets in the BM of pediatric AML patients at diagnosis (Dx) and non-leukemic controls (Mann-Whitney test). (D) Comparison of the time intervals between diagnosis-EOI1 and EOI1-EOI2 (Mann-Whitney test).

**Supplementary Figure 2.**
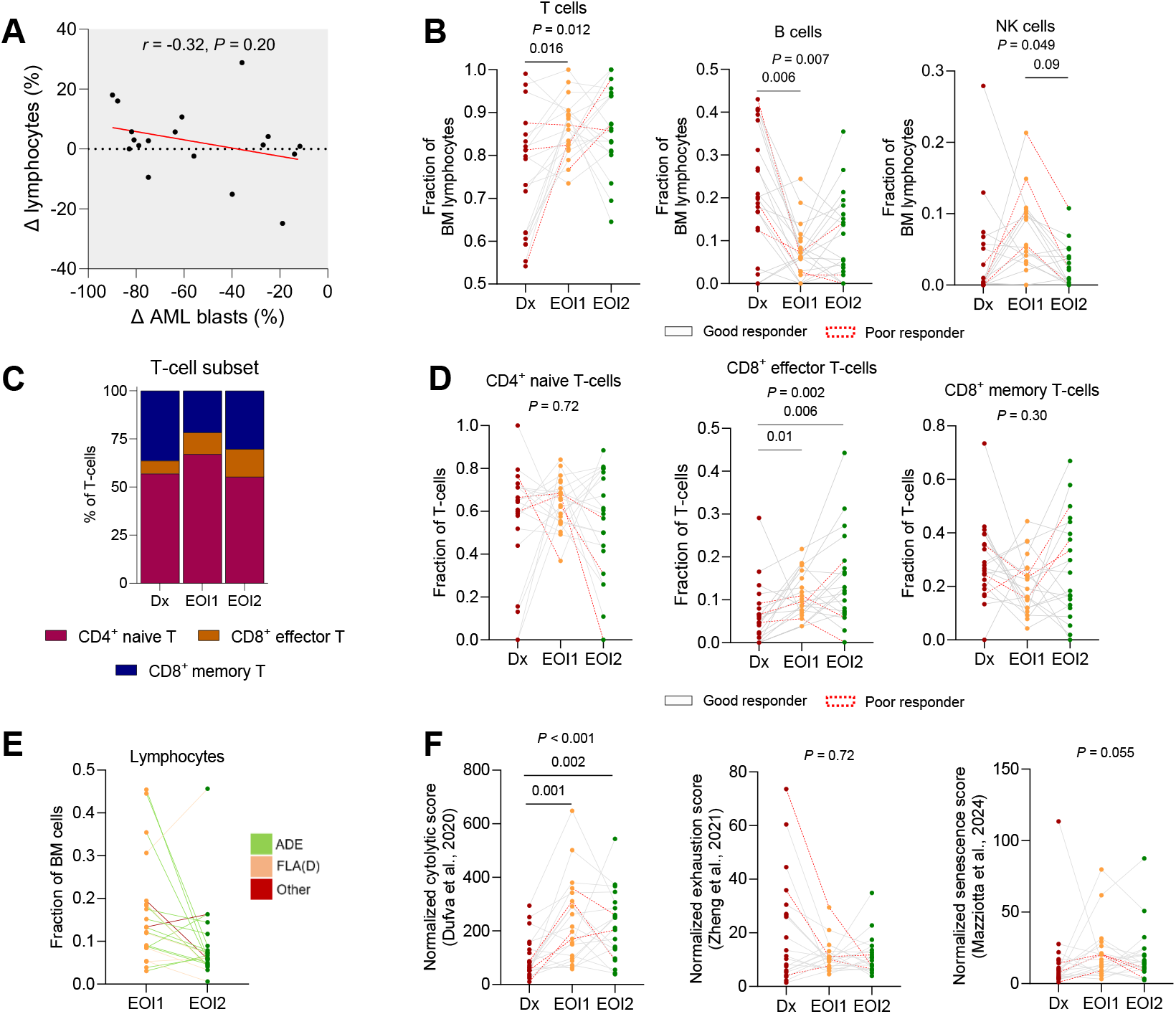
Bone marrow lymphocyte and gene-expression based T-cell functionality dynamics in the primary study cohort. (A) Correlation between the change in AML blasts (measured by flow cytometry) and change lymphocytes between diagnosis and EOI1, calculated using Pearson correlation. (B) Comparison of the fraction of T-, B-, and NK-cells out of BM-lymphocytes throughout induction therapy (Friedman test followed by Dunn’s multiple comparison test; the top value corresponds to the Friedman test, and the lower values indicate Dunn’s test results). (C-D) Fraction of T-cell subsets out of T-cells throughout induction therapy, on average (C) and for each patient (D; Friedman test followed by Dunn’s multiple comparison test). (E) Change in lymphocyte abundance out of BM cells from EOI1 to EOI2, colored according to received therapy during induction 2. (F) Comparison of gene-expression based T-cell functionality scores throughout induction therapy (Friedman test followed by Dunn’s multiple comparison test). Genes for each score: *GZMA, GZMH, GZMM, PRF1*, and *GNLY* (cytolytic score), *PDCD1, TOX, CXCL13, TIGIT, CTLA4, TNFRSF9, HAVCR2, LAG3* (exhaustion score), *NKG7, TBX21, CD38, FCRL6, FCGR3A, C1orf21, PRF1, ENO1, GNLY, ZEB2, CX3CR1, FGFBP2, KLRD1, GZMB, ZNF683, CD226, GZMH, ADGRG1, PLEK* (senescence score). All scores have been normalized for the CIBERSORTx-based T-cell abundance.

**Supplementary Figure 3.**
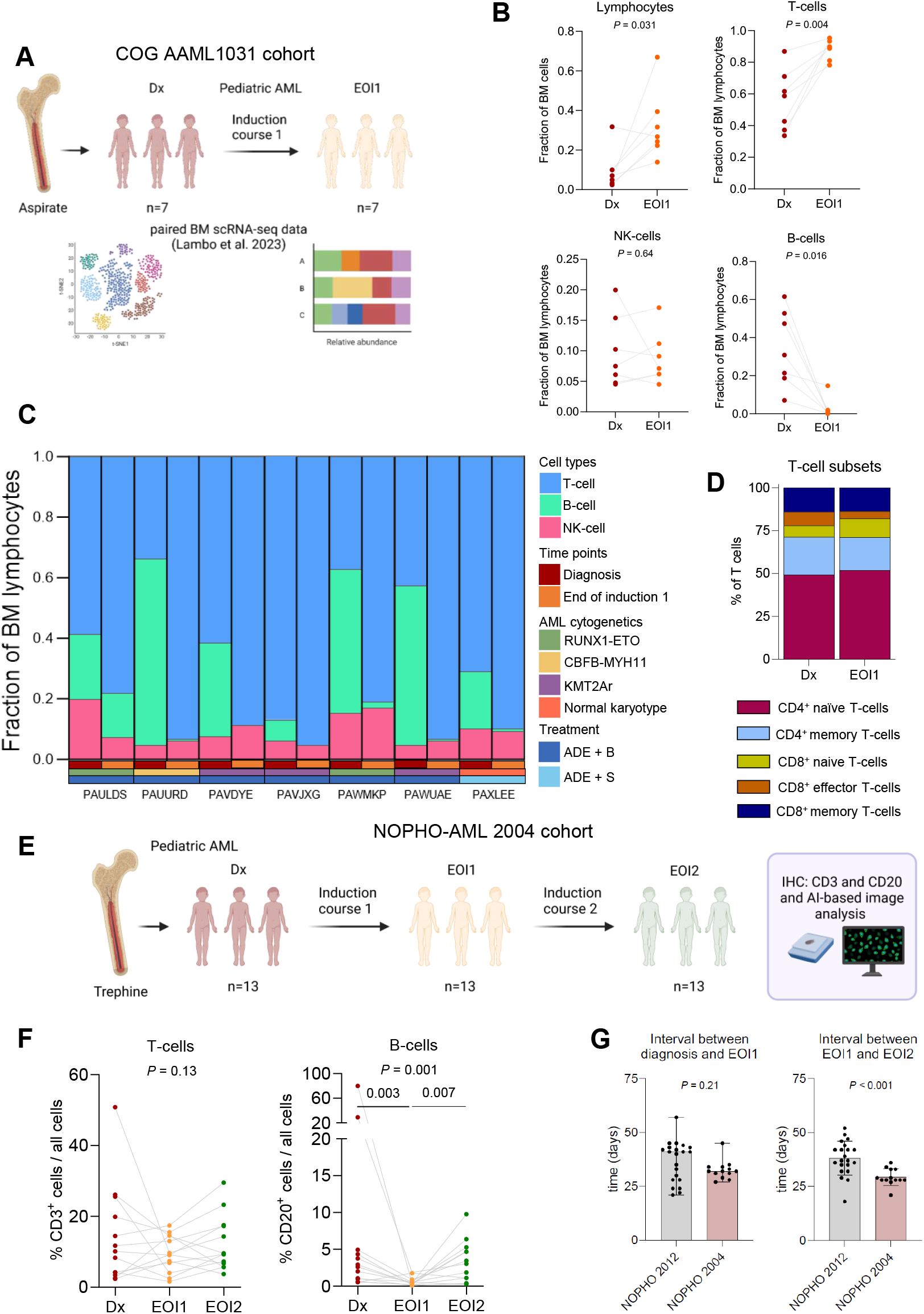
Bone marrow lymphocyte dynamics across international pAML treatment protocols. (A) Schematic overview of the study population in the COG (Children’s Oncology Group) cohort. Dx: diagnosis; EOI1: end of induction 1; BM: bone marrow; scRNA-seq; single-cell RNA-sequencing. (B) Comparison of the scRNA-seq-based fractions (relative abundances) of lymphocytes, T-, B-, and NK-cells at Dx and EOI1 (Paired t test in case of normally distributed data (T- and NK-cells), Wilcoxon matched-pairs signed rank test for non-normally distributed data (lymphocytes and B-cells). (C) Overview of the fraction of lymphocyte populations at diagnosis and end of induction 1, derived from scRNA-sequencing data. ADE: cytarabine, daunorubicin, and etoposide; B: bortezomib; S: sorafenib. (D) Overview of the fractions of T-cell subsets relative to all T-cells at diagnosis and end of induction 1. (E) Schematic overview of the study population in the NOPHO-AML 2004 cohort. IHC: immunohistochemistry; AI: artificial intelligence. (F) Comparison of the CD3^+^ T- and CD20^+^ B-cell abundance from diagnosis throughout induction therapy (Friedman test followed by Dunn’s multiple comparison test; the top value corresponds to the Friedman test, and the lower values indicate Dunn’s test results). (G) Comparison of the intervals (days) between diagnosis-EOI1 and EOI1-EOI2 for the NOPHO 2012 and NOPHO 2004 protocols (Mann-Whitney test).

## Supplementary Tables

**Supplementary Table 1. Characteristics of patients in the primary study cohort**.

**Supplementary Table 2. Characteristics of patients in the COG cohort**.

**Supplementary Table 3. Characteristics of patients in the NOPHO-AML 2004 cohort**.

**Supplementary Table 4. Antibodies used in the *ex vivo* cytotoxicity assays**.

**Supplementary Table 5. Antibodies used for immunohistochemistry**.

## Notes

### Competing Interest Statement

O.H. receives institutional research support from Syndax and Roche. CMZ receives institutional research support from Pfizer, Abbvie, Takeda, Jazz, Kura Oncology, Gilead, and Daiichi Sankyo, provides consultancy services for Kura Oncology, Bristol-Myers-Squibb, Novartis, Gilead, Incyte, Beigene, and Syndax, and serves on advisory committees for Novartis, Sanofi, and Incyte. The remaining authors declare no competing financial interests.

