## Supplementary Data for "Bone marrow lymphocyte dynamics during chemotherapy in pediatric acute myeloid leukemia"

to

Koedijk et al.

**Supplementary Methods**

**Ethical regulations**

Pediatric AML bone marrow (BM) biopsies (trephines) obtained at Aarhus University Hospital (Aarhus, Denmark) were leftover material from standard of care procedures, and all patients and/or their guardians provided written consent for the research use of leftover specimens. The publicly available single-cell RNA-sequencing data utilized in this study was acquired for a study approved by the Fred Hutchinson Cancer Research Centre and the Children’s Oncology Group (COG) Myeloid Disease Biology Committee, as described previously^1^.

**Bulk RNA-sequencing data**

BM bulk RNA-sequencing (bulk RNA-seq) data from pediatric AML patients at diagnosis (n=50) were generated per routine diagnostics and obtained from the Princess Máxima Center Biobank and Data Access Committee. Briefly, total RNA was isolated from BM mononuclear cells (BMMCs), and RNA-seq libraries were generated from 300 ng RNA, and sequenced using the NovaSeq 6000 (2x150 bp; Illumina). After pre-processing, RNA-sequencing typically yielded 60-100 million raw reads, which were then aligned to the GRCh38^2^ reference genome using the gene annotations provided by GENCODE version 31^3^. Control BM bulk RNA-seq data came from non-leukemic samples, including four previously published cases^4^ (children with early-stage rhabdomyosarcoma and no malignant BM infiltration) and three newly generated controls (healthy pediatric siblings acting as a donor for a patient undergoing allo-HCT). In addition, BM bulk RNA-seq was performed on paired samples obtained at specific time points during treatment: end of induction 1 (n=21), end of induction 2 (n=21), end of consolidation 1 (n=6). Each sample was collected right before the start of the subsequent treatment course, with time intervals detailed in **Table S1**. We selected these BM samples to capture the BM in its most regenerated state between chemotherapy courses. For the three patients with a poor response to the first induction course (AML5, AML40, AML47), the subsequent chemotherapy course was initiated immediately upon the detection of ≥5% leukemic blasts. For those with a good response (n=18), hematological regeneration was awaited before proceeding with the next course. Further details on the treatment protocol are provided below.

**Primary study cohort: AML treatment regimen and response criteria**

Acute myeloid leukemia (AML) patients in the primary study cohort were treated according to the NOPHO-DBH AML-2012 protocol^5^. Standard protocol treatment consisted of two induction courses (MEC: mitoxantrone, etoposide, and cytarabine; ADE: cytarabine, daunorubicin, and etoposide) and three consolidation courses (HAM: high-dose cytarabine and mitoxantrone; HA3E: high-dose cytarabine and etoposide; FLA: fludarabine and cytarabine). Initially, induction course 1 consisted of MEC or DEC (daunorubicin-etoposide-cytarabine) and induction course 2 of ADE or FLAD (fludarabine-cytarabine-daunorubicin; randomizations). However, MEC and ADE outperformed the other courses and therefore became the standard treatment as of January 2019^5^ (**Table S1**). Patients in the primary study cohort were diagnosed and treated between July 2019 and November 2024, when the standard induction therapy was MEC and ADE. High-risk patients (those with poor responses to standard therapy or those with a *FLT3*-ITD without a concurrent *NPM1* mutation) received, if feasible, an allogeneic hematopoietic cell transplantation (allo-HCT) after HAM (high-dose cytarabine and mitoxantrone). Standard risk patients with an inv(16) received only two consolidation courses instead of three, omitting HAM. Some treatment adjustments occurred, which are provided in **Table S1**. As per the NOPHO-DBH AML-2012 protocol^5^, bone marrow (BM) aspirates were obtained at day 22 from the start of course one. Patients with ≥5% leukemic blasts in the BM after course one (=poor response to induction one) proceeded immediately to the second course. Accordingly, the day 22 BM aspirates in patients with a poor response to the first induction course one were the “end of induction one” (EOI1) samples. In case of <5% leukemic blasts (=good response to induction one), patients awaited hematological regeneration before starting the second course, with weekly BM aspirates conducted in the interim. In these patients, the BM aspirate obtained right before the start of the subsequent course was referred to as EOI1. Patients with ≥5% leukemic blasts in the BM after course one were subsequently evaluated at day 22 after course two, whereas patients with <5% leukemic blasts after course one were evaluated just before the start of consolidation treatment. EOI2 samples were classified using the same criteria as applied to EOI1 samples. Beyond EOI2, BM samples were obtained only right before the start of the subsequent course (upon hematological recovery; end of consolidation one). Complete remission (CR) was defined as <5% leukemic cells in the BM, as determined by either morphology or flow cytometry, with clear evidence of hematological regeneration (ANC ≥ 0.5 x 10^9^/L and platelets ≥ 50 x 10^9^/L) and no signs of leukemia elsewhere. Refractory disease was defined as the presence of ≥5% leukemic cells in the BM after the second induction course. Whereas patients in CR continued the protocol according to their risk-stratification, patients with refractory disease received salvage therapy.

**Single-cell RNA-seq-guided immune deconvolution**

To create a reference for CIBERTSORTx^6^ (a so-called signature matrix) based on healthy BM cells, we used healthy BM single-cell RNA-seq data from previously published work (Granja et al.^7^). In short, single cells from three healthy BM samples were analysed and filtered to exclude cells with over 15% mitochondrial reads, fewer than 1000 unique molecular identifiers, and fewer than 500 expressed genes. After merging similar cell types and excluding ribosomal genes, we randomly selected 1000 cells per cell type and uploaded them to the CIBERSORTx platform (https://cibersortx.stanford.edu/). Accordingly, the signature matrix was created using the following settings: quantile normalization enabled, kappa of 999, q-value of 0.01, 300-500 barcode genes, minimal expression of 1, three replicates, sampling at 0.5, and no exclusion of non-hematopoietic genes. To assess the accuracy of immune deconvolution using CIBERSORTx with this healthy BM reference, we utilized previously generated scRNA-seq from 27 pediatric AML cases at various disease stages (diagnosis, remission, and/or relapse^1^). Pseudo-bulk profiles were created in R (version 4.4.1) using Seurat (version 3)^8^. Based on cell type annotations from Granja et al.^7^, cells were grouped within the Seurat object, and their expression data was aggregated into pseudo-bulk profiles using Seurat’s AggregateExpression function. This function sums the expression values of cells, effectively creating a “bulk” tissue-level expression profile per patient per treatment time point. Importantly, pseudo-bulk profiles generated from single-cell RNA-sequencing data have been shown to closely resemble bulk RNA-sequencing data^9^. We then applied immune deconvolution to these pseudo-bulk profiles using CIBERSORTx with the healthy BM reference, without enabling batch correction, quantile normalization, or absolute mode. The deconvoluted estimates were subsequently compared with relative cell type abundances of the original scRNA-seq dataset that was annotated using the same healthy BM reference dataset^7^, as previously described^1^. Following this benchmarking analysis, we applied CIBERSORTx with the healthy BM reference to BM bulk RNA-seq data from our primary study cohort (normalized to counts per million; CPM; both in the absolute and relative cell type abundance modes).

**T-cell diversity analysis**

To analyse the diversity of the T-cell receptor (TCR) repertoire from diagnosis throughout induction therapy, we leveraged MiXCR^10^ (version 4.6.0) to reconstruct the TCR repertoires from BM bulk RNA-seq data. First, BAM files were converted to FASTQ format, and MiXCR was applied to these FASTQ files. After preprocessing, MiXCR aligns raw sequencing reads against reference databases of V-, D-, J-, and C segments. It then identifies reads spanning the complementary determining region 3 (CDR3) and assembles overlapping reads into CDR3 sequences. To study unique and identical TCR-β CDR3 sequences, we utilized the respective clones.tsv files generated by MiXCR, where each distinct CDR3 amino acid sequence was denoted as a unique clone. Clones with large insertions or deletions (shown as “_”) or premature stop codons (shown as “*”) were classified as non-functional. Importantly, we only considered clonotypes detected by at least two reads, and calculated the T-cell diversity using the Shannon Diversity Index (SDI)^11^:

$$H =\sum_{i=1}^{n} [\left( pi \right) x\ln\left( pi \right)]$$

*pi:* frequency of clonotype *i.*

*n:* number of unique clonotypes in the sample.

To allow for a comparison of the TCR-β CDR3 diversity between samples with different frequencies of T-cells, we normalized the SDI to the number of unique clones (log10(unique clones + pseudo-count of 1)).

***Ex vivo* assays**

To investigate the cytotoxic capacity of T-cells in between chemotherapy courses, AMV564 (BioSource, Cat# MBS1563606), a bispecific T-cell engager targeting CD33 and CD3 was used. Since all patients with available samples for *ex vivo* testing had hardly any blasts remaining after induction therapy (<1% BM blasts at EOI1/EOI2; **Table S1**), we co-cultured EOI1/EOI2 samples (including CD3^+^ T-cells) with CD33^+^ myeloid cells derived from CD3^+^ T-cell depleted (EasySep™ Human TCR Alpha/Beta Depletion Kit (StemCell Technologies)) diagnostic BM samples for three days (effector-to-target (E:T) ratio of 1:3; all CD33^+^ cells were regarded as ‘targets’), in the presence or absence of AMV564 (50 ng/mL). A 1:1 mix of AML medium and T-cell medium was used for co-culture assays. AML medium consisted of SFEMII (StemCell Tech), SCF (150 ng/mL), TPO (100 ng/mL), SR1 (750 nM), UM171 (135 nM), IL-3 (10 ng/mL) and FLT-3 (10 ng/mL; all purchased from PeproTech) and 100 μg/mL Primocin (InvivoGen). The T-cell medium contained RPMI + Glutamax (Gibco), 10% human male serum (Sigma-Aldrich), IL-2 (100U/mL), IL15 (10 ng/mL) and IL7 (10 ng/mL) purchased from PeproTech plus 100 μg/mL Primocin. After 3 days, the remaining viable CD33^+^- and T-cells were stained and counted with flow cytometry (CytoFlex LX and CytoFlex S, Beckman Coulter) using corresponding antibodies (**provided in Table S4**)**.** Specific lysis induced by AMV564 was calculated as =

100 - [(Number of viable CD45dim or CD45^+^CD33^+^ cells in treated / non-treated conditions) *100].

To further evaluate T-cell functionality, we measured changes in the T-cell activation markers CD25 and CD137, granzyme B expression on the T-cell-fraction of the co-cultures treated with or without 50 ng/mL AMV564. To do so, after 3 days of co-culture, cells were treated with 1x monensin (Biolegend) for 4 hours at 37°C followed by staining with surface markers (antibodies listed in **Table S4)**. Cells were then fixed for 30 minutes (CytoFix, BD BioSciences) at 4°C, and permeabilized (Phosflow™ Perm Buffer III, BD BioSciences) for 30 minutes on ice. Staining with 24ug/mL of granzyme B or isotype control was proceeded on ice before acquiring samples with flow cytometry. Moreover, we evaluated T-cell numbers and fractions of subsets (naive, effector memory, central memory, TEMRA) after the 3-day co-culture (antibodies listed in **Table S4**).

**COG AAML1031 cohort**

The previously published dataset (Lambo et al., 2023^1^) utilized to perform benchmarking analyses of the healthy BM reference included scRNA-seq from fourteen paired diagnostic and EOI1 BM samples from seven treatment-naïve pediatric AML patients. These scRNA-seq data were annotated using the healthy BM scRNA-seq dataset by Granja et al.^7^ and allowed for comparative analyses between diagnosis and EOI1. Albeit scRNA-seq data for matched diagnosis and EOI2 BM samples were available for six patients, the lack of EOI1 data for these patients precluded investigations into how treatment according to the AAML1031 trial^12^ affected the abundance of BM lymphocytes throughout induction therapy. As denoted in the main text, six patients received ADE and bortezomib, while one patient was treated with ADE and sorafenib because of *FLT3*-ITD-positivity. Patient characteristics are provided in **Table S2**.

**NOPHO-AML 2004 cohort (Aarhus University Hospital)**

We previously generated single-stain immunohistochemistry (IHC: CD3 and CD20) data for diagnostic formalin-fixed and paraffin-embedded BM biopsies (trephines)^13^ from pediatric AML patients treated at the Aarhus University Hospital. For thirteen patients, consecutive samples at EOI1 and EOI2 were also available (n=26 additional samples), on which we performed conventional IHC using antibodies against CD3 and CD20, as described previously^13^. Briefly, IHC was applied to consecutive 4 μm sections employing a Ventana Benchmark Ultra (Roche, Basel, Switzerland) automated staining instrument according to manufacturers’ instructions (antibodies and suppliers are provided in **Table S5**), and subsequently scanned using a NanoZoomer scanner (Hamamatsu, Shizuoka, Japan). Whole-slide digital image analysis was done in QuPath^14^, utilizing the deep learning-based cell segmentation tool StarDist^15^ and the machine learning-based Random Trees classifier. CD3- and CD20 cell type abundances were quantified as percentage of all nucleated cells to allow for a direct comparison between diagnostic (low amount of faT-cells) and EOI1/EOI2 samples (high amount of faT-cells, occupying a relatively larger area of the BM compared to diagnostic samples with overt AML). All thirteen patients were treated using AIET (cytarabine, idarubicin, etoposide, and 6-thioguanine) as the first induction course and AM (cytarabine and mitoxantrone) as the second^16^. Patient characteristics are provided in **Table S3**.

**Statistical analysis**

All statistical analyses were conducted using GraphPad Prism v10.0.2 (GraphPad Software, LA Jolla, CA, USA). For data that did not follow a normal distribution, comparisons between two independent groups were made using the Mann-Whitney test, while paired groups were analysed with the Wilcoxon matched-pairs signed rank test. For data following a normal distribution, unpaired and paired t tests were used accordingly. To assess correlations between two variables, Pearson rank correlation coefficient (Pearson’s *r*) was applied. For multiple comparisons where residuals did not follow a normal distribution, we used the Friedman test followed by Dunn’s multiple comparison test with Bonferroni correction. When multiple P-values are reported in figures, the top value corresponds to the Friedman test, and the lower values indicate Dunn’s test results. A P*-*value of less than 0.05 was considered statistically significant for all tests.
